# Elasto-Osmotic Phase Separation in Confluent Cellular Tissues

**DOI:** 10.64898/2026.05.29.727481

**Authors:** Jasper J. Michels

## Abstract

Biomolecular condensates that form via liquid-liquid phase separation (LLPS) of, most prominently, intrinsically disordered proteins (IDPs) are ubiquitous in eukaryotic cells and responsible for regulating a plethora of biological functions. Amongst these, they contribute to regulating cell motility, either individually within an extracellular matrix or collectively within confluent epithelial tissue. In this computational study we focus on the latter with the aim of investigating whether the mutual exertion of mechanical forces during collective migration in an epithelium can principally trigger cytoplasmatic LLPS. Since present models for confluent epithelial motility have so far only considered cells that are devoid of phase separating (protein) solutes, we extend a common multiphase approach for 2D cell motility with a mixing contribution including any number of protein solutes. Our model considers the phase behavior in both intracellular and extracellular regions and determines to what extend the membrane is permeated by the solutes under the influence of mechanical and osmotic forces. Our initial calculations unlock a very rich behavior involving formation and dissolution of condensates during migration, as well as an impact of LLPS on the very nature of the motility itself, through feedback mechanisms which may bear biological relevance.

## I. INTRODUCTION

Epithelium is a thin layer of barrier tissue that lines and protects surfaces of organs, body cavities, blood vessels and skin. Epithelial cells are typically motile: they migrate driven by actin (de)polymerization and myosin action in combination with transient formation of focal adhesions at the underlying substrate [1]. Being eukaryotic cells, epithelial cells contain phase separated droplets of compartments referred to as ‘biomolecular condensates’, that form via liquid-liquid phase separation (LLPS) and serve a plethora of biological functions [2, 3]. Amongst these functions, we also find the regulation of cellular and tissue dynamics and mechanics. For instance, biomolecular condensates have been shown to regulate actin polymerization [4], cytoskeletal integrity [5], assemble epithelial and tight junctions [6, 7], regulate focal adhesion dynamics [8] and spatially organize membrane receptors [9].

Many biomolecular condensates contain a large fraction of intrinsically disordered proteins (IDPs), which bear domains that are uncapable of folding into stable secondary structure. For that reason, IDPs are generally hydrophobic and, as it seems, near their saturation point to allow for the cell to trigger condensation or de-condensation in response to relatively subtle stimuli, such as changes in pH, salt, temperature or chemical modification. Also mechanical cues are being recognized as direct or indirect means of inducing biocondensation [8, 10, 11, 12] and regulators for condensate dynamics [13]. Conversely, biomolecular condensates contribute to the regulation of cellular mechanical properties, including cytoskeletal stiffness and cell adhesion, playing key roles in mechano-transduction and the cell’s response to mechanical stimuli [13].

In regard of the above, if the mean concentration of scaffolds for biomolecular condensates indeed typically resides near a phase boundary, we propose that it is not too much of a stretch to imagine that small changes in cell volume can lead to a crossing of the phase boundary, thereby inducing LLPS, of course given the cell membranes are semipermeable and in response to compression and extension mostly allow for transport of water and perhaps small ions. In confluent tissues, small changes in cell volume may occur under influence of mechanical stresses that arise as a result of cell motion [14]. Although detailed experimental evidence for such a trigger for condensate formation still has to emerge, the implications are potentially far-reaching as it would provide for a means for regulating long-range biological function based on (sub-)cellular mechanisms.

In this work we computationally explore such motility-triggered condensate formation in confluent 2D tissues, within a context we refer to as ‘osmo-elastic’ phase separation on account of the fact that the observed phenomenology arises from a coupling between elastic and osmotic forces. Our aim is not to irrefutably prove the occurrence of the mechanism in biology, but to provide a sense of plausibility by demonstrating and detailing the mechanisms underlying the process, as well as identifying feedback mechanisms and their physical implications.

Computational tools for investigating osmo-elastic phase separation in confluent epithelia do not exist. Well-known multiphase-field models for cellular motion, whether individually in an extra-cellular matrix or in confluent ensembles are available [15, 16, 17], but only consider cells that are devoid of (phase separating) solutes. For that reason, we extend a minimal version of such a model with a mixing contribution that takes into account any number of solutes that have interaction parameters evaluated for the intra- and extracellular regions separately. The resulting multiphase-multicomponent describes coupled phenomena between motility and phase separation, driven by elastic changes in cell size and shape and as a function of the partitioning strength of the solutes across the cell membranes.

The remainder of this work is organized as follows: in Section II we describe our model in detail, placing emphasis the extension of the multiphase free energy with a multi-component mixing contribution, as well as the consequences for coupled dynamics equations and mass conservation. In Section IIIA we investigate the osmo-elastic phase behavior of a single static cell containing a binary cytoplasm (single solute), both in equilibrium and maintained out of equilibrium by applying an external pressure. We map out pressure-composition phase diagrams and compare full and partial miscibility in the cytosol. In Section IIIB we give a flavor of the extent of the phenomenology by exploring two dynamic scenarios on multi-cell epithelia of confluent motile cells. We show how phase separation is influenced by the characteristics of cellular motion, *i.e*. deterministic or stochastic, and compare osmo-elastic phase behavior as a function of the effective permeability of the cells. Interestingly, we identify feedback mechanisms in which motility-induced condensation affects the fluidity of the epithelium itself. In Sections IV we summarize and discuss our results and provide general conclusions in Section V.

## II. MODEL

Our 2D computational epithelium comprises *n* confluent cells that coexist with an extracellular environment (ECE). We refrain from using the term ‘extracellular matrix’ to avoid overclaiming the resemblance with a real biological tissue, although we stress that the model is extendable to including specific mechanical properties of both the cells and the ECE. A solution containing *m*-1 solutes (*m* indexing the solvent) partitions across the tissue, with its spatiotemporal distribution being determined by affinities specified between all component and in each cell and the ECE individually. A central feature of our model is that these solutes, let’s say proteins, genetic material etc., should be able to phase separate or remain dissolved in either or both the cytoplasm and the ECE, depending on concentration temperature, polarity, etc. We write the free energy for this system as a function of sets of conserved (volume fractions) {*c*_1≤*i*<*m*_(**r**)} and non-conserved order parameters {*φ*_1≤*k*≤*n*_(**r**)} as:

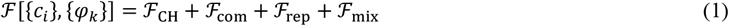

where {*φ*_1≤*k*≤*n*_(**r**)} identify the locus of each cell, assuming values of 1, 0 and 0 < *φ*_*k*_ < 1 in the cytoplasm, in the ECE and at the interface, respectively.

The terms on the RHS of Equation 1 respectively represent a Cahn-Hilliard free energy, a contribution due to cell compressibility, a penalty due to spatial overlap between cells and a mixing free energy. These are defined as:

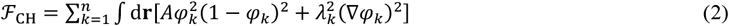

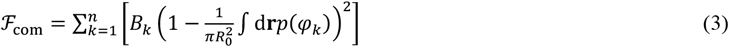

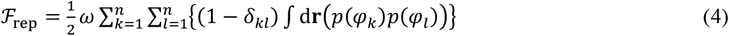

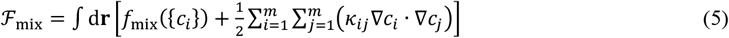

The first three contributions are based on ref. [18], the fourth being the new extension responsible for the phase behavior. The first term in the integrand of Equation 2 represents a Landau-type double well function with height *A*, which guarantees coexistence between a cell and the ECE. The second penalizes order parameter gradients, with gradient energy coefficients 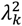. The compressibility Equation 3 holds a soft constraint penalizing the deviation of the cell volume from a ‘target value’ 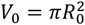, with *B*_*k*_ representing the associated energy scale. The interpolation function

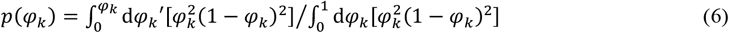

is common in materials science applications of phase field models [19, 20] and assures thermodynamic consistency in that in equilibrium *φ*_*k*_ is zero/unity outside/inside cell and that the condition d_*φk*_ℱ = 0 is met in both the cytoplasm and the ECE. Spatial overlap between order parameter fields is penalized by ω (Equation 4).

As mentioned above, the mixing free energy (Equation 5) is the essential new contribution in this model. Its local contribution is a multicomponent Flory-Huggins-type free energy, given by

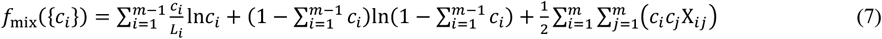

with *L*_*i*_ a scaled molecular size of solute *i* and *L*_*m*_= 1 referring to the solvent. The first two terms on the RHS of Equation 7 are contributions due to translational entropy and excluded volume. The third term is due to pairwise interaction, where the interaction parameters are expressed as

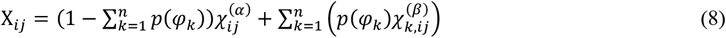

interpolating between extremes 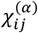 in the ECE and 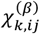 in cell the cytoplasm of cell *k*, assigned to the regions outside and inside a cell, to which we refer as the α- and β-phase. Unlike the cells, the solution is considered incompressible, so 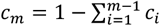. As we shall see below, the magnitude of the interaction parameters and in particular the difference 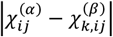 determine to what extent the solute prefers to reside in the cytosol or the ECE and hence, in effect, the ‘(semi-)permeability’ of the cell membrane. Finally, the second term in the integrand of the total mixing free energy (Equation 5) penalizes gradients in solute concentration, with *κ*_*ij*_ the associated gradient energy coefficients. Cross contributions of gradients in concentration and order parameters are neglected for simplicity and computational tractability.

For modeling the spatiotemporal changes in shape, size and velocity of each cell, we use an earlier proposed [21] advection-extended Model A in the Hohenberg-Halperin classification [22]:

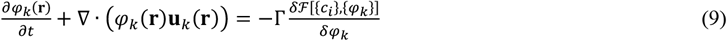

with **u**_*k*_(**r**) the velocity of cell *k* given as a locally resolved vector field and Γ a kinetic coefficient governing the interconversion between cytosol and ECE. Of course, in our particular case changes in the order parameters are coupled to those in the local composition via the free energy. The local cell velocity relates to a force according to [21]:

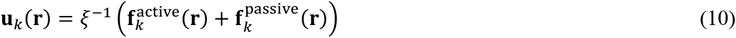

with ξ a friction coefficient and **f**_*k*_ forces acting on cell *k*. The active contribution 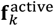 drives cell motion and maintains the system out of equilibrium and 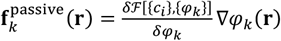 represents a thermodynamic restoring force.

Expressions for 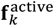 have been developed with the aim of closely modeling the biological mechanisms responsible for cell motion, such as actin (de)polymerization, myosin contraction and inter-cell viscous friction [17, 23]. In view of the present work’s focus on the demonstration of osmo-elastic phase behavior, we do not aim to mimic biological motility as closely as possible, although it would certainly be of interest to refine the model as such in future work. For now, it suffices for 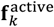 to yield a minimal representation of confluent motility based on deterministic and stochastic contributions that are tunable relative to each other. Further following ref. [21] we define: 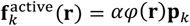, with *α*a coupling parameter and **p**_*k*_ = (cosθ*k* sinθ*k*), with 0 ≤ θ_*k*_ < 2π a randomly sampled angle. As in ref. [24], the angle evolves in time subject purely to stochastic fluctuations according to ∂_*t*_θ_*k*_ = ζ_*k*_, with an amplitude determined by ⟨_*k*_(*t*)ζ_*k*_(*t*′) = = 2*D*_*R*_δ(*t* − *t*′), with *D*_*R*_ a rotational diffusivity. Hence, *α* and *D*_*R*_ modulate the strength of the deterministic and stochastic components of 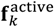. We will perform all calculations at a fixed *α* and vary *D*_*R*_ to investigate the effect of the nature of the motility on the phase behavior of the cytoplasm.

A challenge in obtaining a suitable evolution equation for the solute concentrations is that for solute(s) and solvent to follow cell motion, the change in their local concentration, being a *conserved order parameter*, is subject to a velocity field **u** = ∑_*k*_ **u**_*k*_ obtained based on evolution of the *non-conserved* order parameters {*φ*_*k*_}. As a result, **u** is not necessarily non-divergent, implying that by bluntly coupling *c* to **u** in a convection-diffusion equation, solute might spuriously form or disappear as the cells contract and extend during migration. Applying an equation of state, as is typically necessary to maintain mass conservation when modeling compressible flow, seems inappropriate in this case, as i) pressure is ill-defined in this problem and ii) the mass density of the solution does not depend on cell compression and extension. A suitable ‘equation of state’ would relate the solute concentration to the pressure that a cell locally experiences. However, this is difficult since per Equation 3 ‘cell pressure’ is defined *per cell* rather than locally.

As a simple but effective compromise that nearly perfectly conserves mass and upholds a reasonable coupling between the solute and local (cell) velocity, we decompose **u** in its non-divergent (incompressible) and curl-free components and couple the concentration fields exclusively to the former through the following Model H-type [22] multi-component convection-diffusion equation:

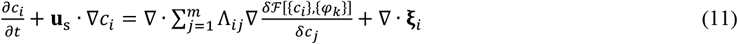

with Λ_*ij*_ mobility coefficients, ξ_*i*_ a stochastic flux due to thermal fluctuations in a multicomponent mixture [25] and **u**_s_ = **u** − ∇𝔇 the solenoidal component of **u**, obtained via a Helmholtz decomposition, with ∇𝔇 representing the curl-free component. We demonstrate below and Section S-II of the Supplementary Information (SI) [26] that applying this approach guarantees near perfect mass conservation, even for highly fluid epithelia, for which cell migration is extensive.

## III. RESULTS

### A. Phase behavior

In order to understand how the model captures the dynamic coupling between cell motion and phase separation in driven, confluent out-of-equilibrium ensembles of *multiple motile* cells, it is instructive to first examine the how the composition and size of a *single stationary* cell (*n*= 1) containing a single solute A (*i.e. m*= 2) are determined by the interplay between osmotic and elastic forces. In effect, what we are interested in is the solute concentration in the ECE and the cell as two coexisting phases, denoted α- and β-phase, as a function of the stiffness of the cell *B*, the solubility of the solute in the two phases and the presence of an external compressive force. We write the local free energy density for each phase *i* ∈ {*α, β*} as:

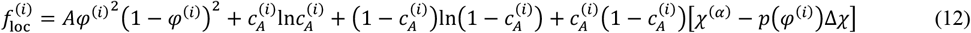

with Δ*X = X*^(*α*)^ − *X*^(*β*)^, *φ*^(*α*)^ = *P*(*φ*^(*α*)^) = 0, *φ*^(*β*)^ = *P(*φ*^(β)^) =* 1 and where, for simplicity, we have taken the relative molecular size of both the solute and the solvent to be *L* = 1. In equilibrium, the osmotic pressure difference between the cell and the ECE is balanced by an elastic potential associated with the cell’s compressibility, or, phrased differently, the resistance of the cell against expansion or compression under influence of osmotic pressure:

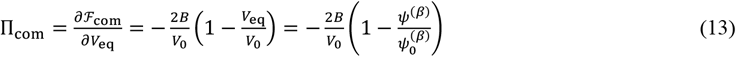

with 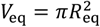 the equilibrium volume of the cell and ψ^(β)^ the cell’s fractional volume within the volume occupied by a ‘unit cell’ containing the cell itself and its characteristic ECE, and 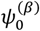 the cell’s fractional volume in the absence of solute partitioning, in which case *V*_eq_ = *V*_0_ and osmo-elastic stresses are absent except for interfacial tension due to gradients in *φ*, which we neglect in the present exercise. The ‘lever rule’ relates ψ^(β)^ to the mean solute concentration 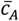 in the unit cell, as well as the solute concentrations in the coexisting phases as 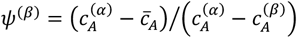.

As ψ^(β)^ enters the expression for Π_com_, in what follows we refer to the latter as the ‘osmo-elastic potential’.

For given *B, V*_0_ and 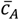, the equilibrium composition of the cell and its ECE, together with the size of the former can be obtained by solving the conditions

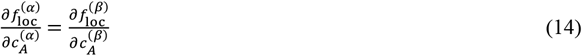

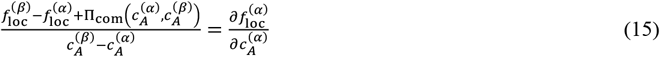

for 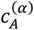 and 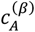. Performing this common tangent construction for a range in any of the control parameters 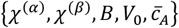 in principle accesses a variety of phase diagrams that demonstrate how cell size and composition is interrelated. Alternatively, one may also calculate the phase diagram bypassing the independent variables by enforcing Π_com_ as a control parameter, *i.e*. treating it as a result of an external mechanical force that maintains the system out of equilibrium, in which case, one should consider *V*_eq_ as a steady-state value. These approaches give complementary insight, which we demonstrate below.

FIG 1 expresses the phase behavior if the solute is partially miscible in the ECE but fully miscible in the cytosol, by defining: *χ*^(*α*)^ = 3, *χ*^(*β*)^ = 1.5. To maintain biological relevance, we consider a low overall solute concentration to guarantee 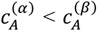. Hence, we do not investigate phase separation in the ECE but rather exploit the form of its mixing free energy to ensure a low 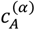, as well as a low to modest dependence on the osmo-elastic potential Π_com_. FIG 1a displays the phase diagram in terms of the variation in the binodal compositions with Π_com_. The common tangents in FIG 1b show how the coexisting (binodal) concentrations relate to the free energies and that Π_com_ is represented by the offset between the free energies of the α- and β-phase. Increasing its value from zero, *i.e*. following the numbering of the common tangents, results in an increase as well as a divergence of the binodal compositions 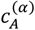 and 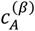. If the magnitude of Π_com_ exceeds the value 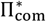, for which the composition of the α-phase coincides with 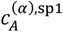 (vertical red dashed line in FIG 1a and 1b), a common tangent no longer exists.

**FIG 1.**
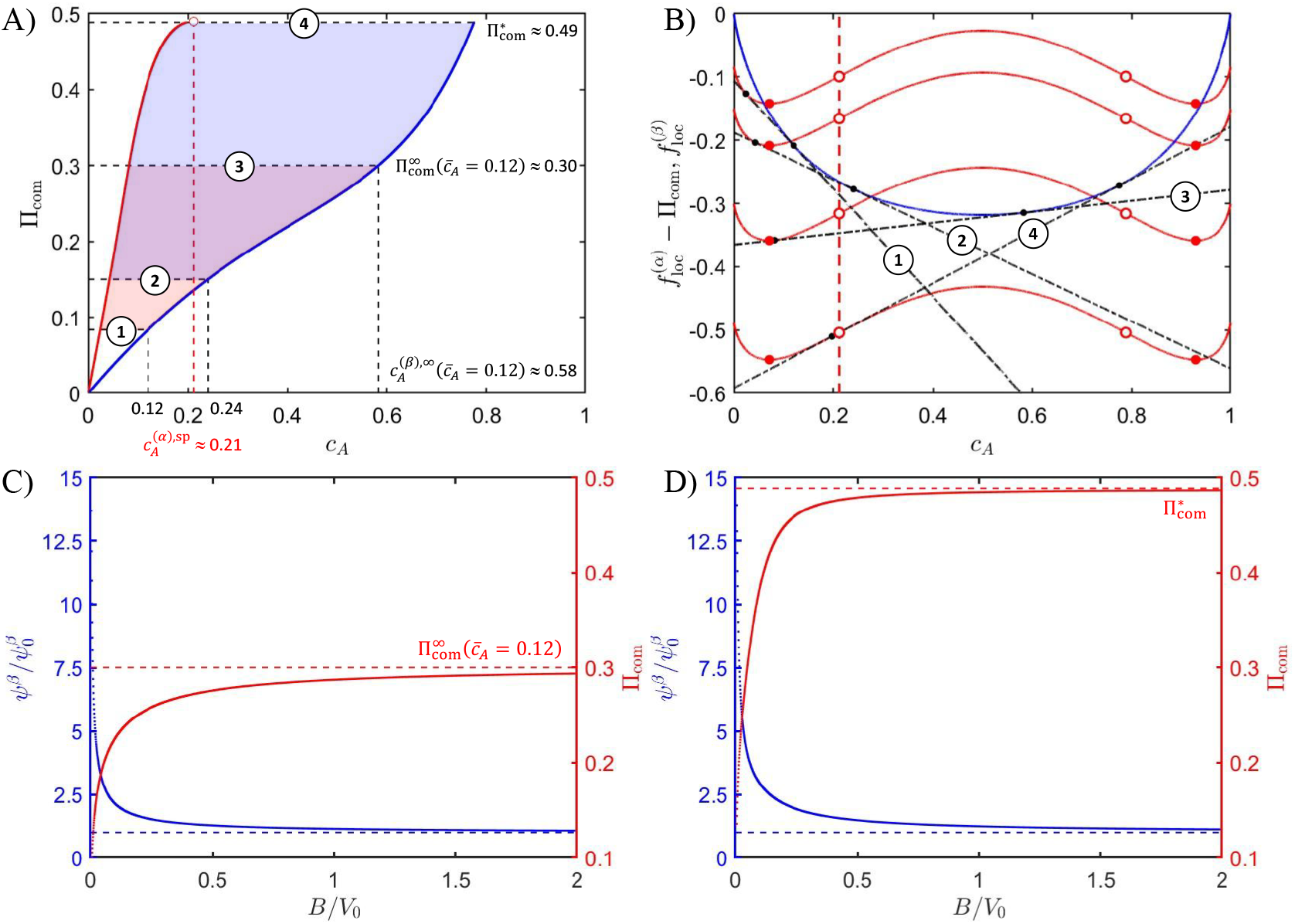
Osmo-elastic phase behavior of a single cell containing a miscible solute. (a) Isothermal phase diagram as a function of the elastic potential with coexisting compositions of the cytosol and the ECE indicated in blue and red. The open red symbol corresponds to the lowest spinodal composition in the ECE. The shaded red and blue areas represent the sections of phase space addressed for 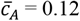 and 0.24, respectively, obtained by calculating the solute partitioning as a function of cell stiffness B rather than Π_com_. (b) Common tangents (dash-dotted) to the mixing free energies of the cell (blue) and the ECE (red), numbered as indicated in panel (a). The mixing free energy of the ECE is off-set by Π_com_. Solid and open symbols respectively represent binodal and spinodal compositions. (c) und (d) Normalized cell size (blue) and elastic potential (red) plotted as a function of the normalized cell stiffness for 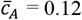 and 0.24, respectively.

Calculating the binodal compositions under a passively varying Π_com_ based on changing the stiffness *B* as an independent variable, reveals something interesting when comparing mean compositions 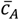 above and below 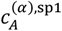.

In case we assume 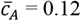, which is smaller than 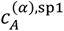, the potential Π_com_ saturates at an asymptotic value 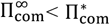 (red curve in FIG 1c). The blue curve in FIG 1c shows that concomitantly the normalized volume of the β-phase approaches unity, implying that the cell volume approaches the target volume *V*_0_, resisting deformation due to osmotic forces. As a result, 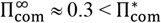 demonstrating that for 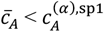 only part of the total phase diagram can be mapped (red shaded region in FIG 1a). In contrast, FIG 1d shows that for 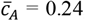, *i.e*. exceeding 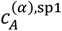, 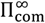 coincides with the limiting value 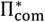, implying that now the binodal compositions can be obtained up to the point where a common tangent ceases to exist (blue shaded region in FIG 1a).

A very different behavior is obtained in the more interesting case for which in *both* the ECE and the cytosol the solute is partially miscible. This scenario is laid out in FIG 2 and forms the bases for the dynamic calculations below. We achieve partial miscibility through: *χ*^(*α*)^ = 4, *χ*^(*β*)^ = 2.5. Again, the A-panel shows the phase diagram, with numbered tie-lines corresponding to the common tangents in the B-panel, but now plotted on a logarithmic concentration scale to reveal intricacies in the behavior of the composition of the ECE. In contrast to the miscible case, now typically *multiple common tangents* can be drawn for 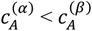, owing to the double-well nature of the mixing free energy of the cytosol (blue curve in FIG 2b). In the phase diagram in FIG 2a we express the consequence of this by plotting corresponding branches in different shades. Whether these compositions physically coexist from a thermodynamic perspective depends on how they relate to the individual binodal and spinodal loci of the individual free energies (vertical red and blue dashed lines in FIG 2a and 2b).

**FIG 2.**
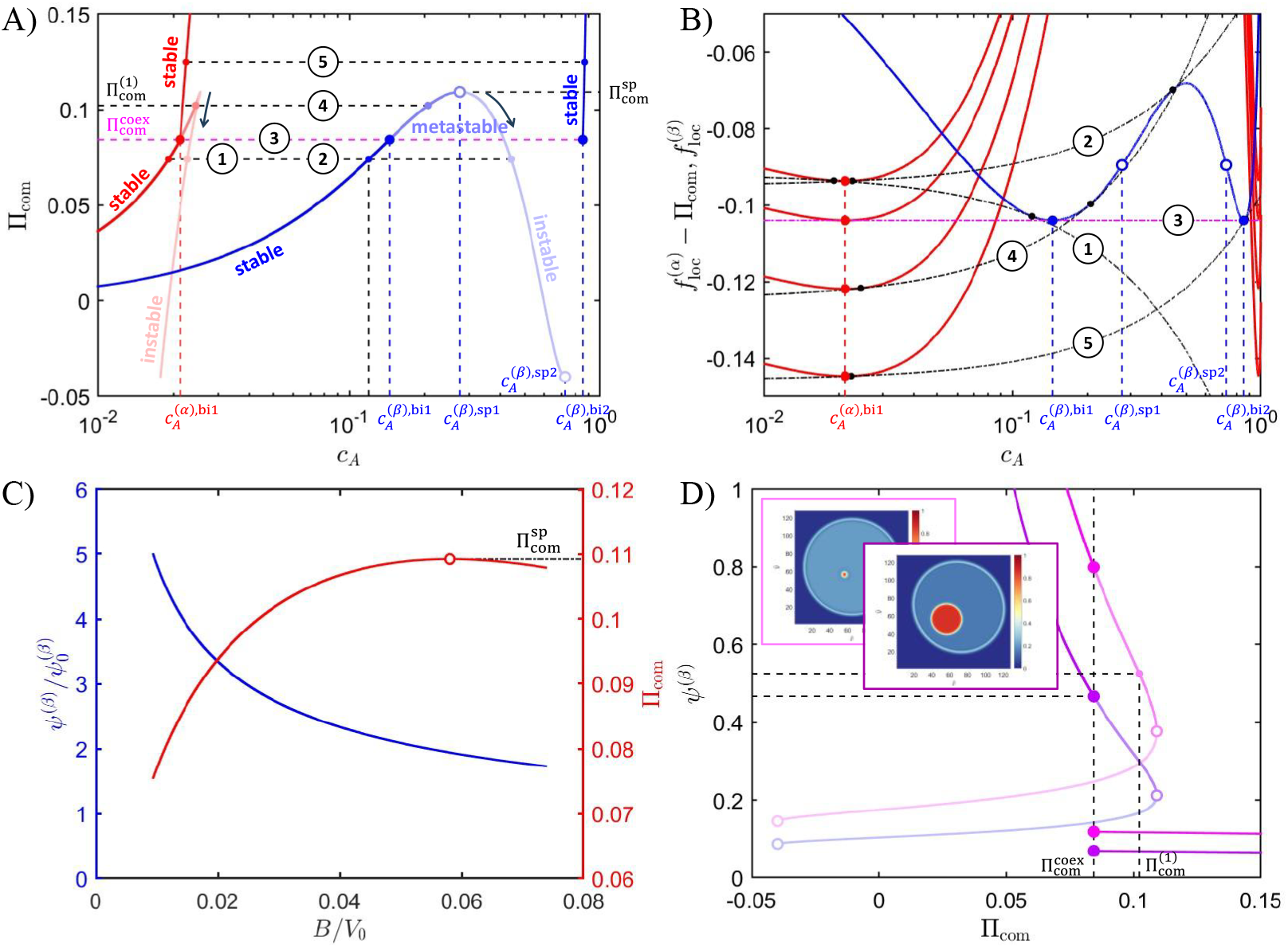
Osmo-elastic properties of a single cell containing a partially miscible solute. (a) Isothermal phase diagram with coexisting compositions of the cell and its ECE indicated in blue and red; stable, metastable and unstable equilibria in dark, medium and light shades. The dashed black lines are tie-lines. (b) Common tangents (dash-dotted) to the mixing free energies of the cell (blue) and the ECE (red), numbered as indicated in panel (a). In (a) and (b), individual binodal and spinodal compositions are indicated by (large) closed and open symbols and vertical dashed lines. (c) Normalized relative cell size (blue) and elastic pressure (red) plotted as a function of the normalized cell stiffness for 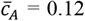. (d) Relative cell size plotted as a function of the elastic pressure. The magenta and purple curves correspond to 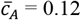 and 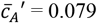. Solid and open symbols represent binodal and spinodal points. Insets (top and bottom): initial and final snapshot of nucleation and growth of a condensate inside a cell. The horizontal dashed lines mark the cell size in the metastable (initial) and stable (final) states.

We relate the shading of corresponding branches of the phase diagram based on whether a specific scenario leads to thermodynamic stability (dark shade), meta-stability (medium shade) or instability (light shade) of the *cytosol*. This convention gives rise to two stable regimes: 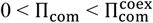 and 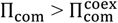, as indicated in FIG 2a, where 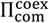 signifies the potential where all four individual binodal compositions fall on a common tangent (see magenta tie line **3**). The transition between these regimes is discontinuous, exhibiting a jump in binodal composition of the cytoplasm. Close comparison between common tangents **1** and **5** reveals that for stable regime i) both coexisting compositions are on the individually stable section of both free energies (so,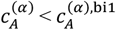 and 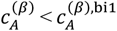,) whereas for stable regime ii) the ECE composition is just metastable 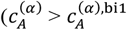.

Furthermore, stable regimes i) and ii) partly coincide with an unstable regime determined by tangents that connect either the individually stable 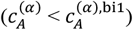 or metastable part 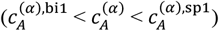 of the ECE free energy with the unstable section of the free energy of the cytoplasm 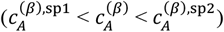. Tie line **2** exemplifies this regime.

For an osmo-elastic potential in the range 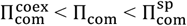, we identify a metastable regime, exemplified by tie-line **4**, whereas for 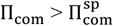 we exclusively identify stable regime ii) with 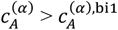 as discussed above. The regime that tie-line **4** is part of is of central importance to the dynamic phenomena associated with osmo-elastic phase separation. In the next section, we show that moving in and out of this regime via motility-induced variations in Πcom causes actual nucleation and dissolution of condensates.

As motility-induced (reversible) condensation in the cytoplasm besides cell compression and extension due to external forces also depends on cell stiffness, we inspect in FIG 2c how Π_com_ is modulated by increasing the stiffness parameter

*B*. The behavior is initially qualitatively similar to the case of a miscible cytoplasm where the elastic potential Π_com_ increases and the size of the cell decreases, leading to a non-linear approach of the ratio 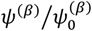 to unity. However, the behavior of Π_com_ is eventually different. Rather than saturating, its value reaches a maximum at 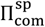 once the contact point of the common tangent at the cytosol free energy reaches the spinodal concentration 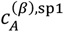.

Mathematically, this is a consequence of the fact that for a common tangent to exist beyond this point requires a reduction in Π_com_, as indicated in FIG 2a by the dark arrows along the light red and blue shaded branches delineating the regime in which the cytosol is thermodynamically instable.

FIG 2d and inset express the changes to a cell brought about by nucleation and growth of a condensate, converting a meta-stable cell characterized by tie-line **4**, osmo-elastic potential 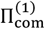 and 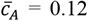 into a three-phase stable equilibrium associated with a pressure 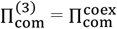, as also indicated in panel (a). The magenta line in panels (a) and (b) (and corresponding closed symbols) indicate the concentrations of the three coexisting phases: one outside the cell: 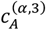 and two inside the cell: dilute phase concentration 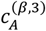 and condensate concentration 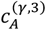. Interestingly, despite the decrease in Π_com_ during condensation, *i.e*. 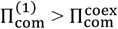, the size of the cell decreases as well. This is due to the fact that the solute concentration in the majority phase 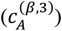 decreases relative to the initial concentration in the cell 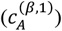, whereas that of the ECE remains virtually unchanged: 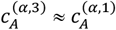, giving rise to the decrease in ψ (see panel (d)), as obtained by Equation 13 for ψ and given *B* and domain size *N*. In effect, although mass is conserved, *i.e*. 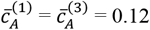, the size of the cell in the stable equilibrium is reproduced in a miscible scenario by omitting the excess solute in the condensate and instead considering an effective new mean concentration 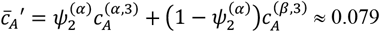 (see caption FIG 2).

### B. Confluent dynamics

In this section we inspect the spatiotemporal non-equilibrium behavior of a confluent epithelium comprising single solute cells by numerically integrating evolution Equations 9 – 11, while varying a specific set of input parameters. All parameters are treated dimensionless and are listed in Table S-I in Section S-II of the SI [26]. In the main text we only mention specific parameters of interest to the presented exercises. All calculations start from a near-equilibrated confluent epithelium, which we generate via nucleation and growth [27], whereby we ensure the target radius of each cell to be sufficiently large to guarantee confluency. We achieve this by defining 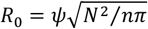, with *N* = 128 the size of the square numerical grid, *n*= 12 the number of cells and ψ = 0.99 a parameter that tunes the confluency. For these settings, the volume occupied by *n*equilibrated cells is larger than the available space determined by the computational box, meaning the cells are generally deformed and compressed relative to their equilibrium volume. Consequently, in the epithelium the confluency decisively contributes to the osmo-elastic potential experienced by a motile cell through its extension and compression resulting from the mechanical coupling with its neighbors. During these processes composition gradients develop and a cell exchanges material, *i.e*. solute and solvent, with the ECE and neighboring cells. The exchange of solvent trivially follows from the global incompressibility constraint: if a cell extends it takes up solvent and releases it upon compression. The exchange of the solute is subtly different. If there is no preference of the solute to reside in either the ECE or the cytosol, the cells effectively behave as if there were fully permeable and the solute will be exchanged much the same as the solvent. However, if the solute partitions in favor of either of the two phases, as determined by *χ*^(*α*)^ and *χ*^(*β*)^, the cell behaves as if its ‘membrane’ were, in a thermodynamic sense, ‘semipermeable’. Importantly, in this case the loss or gain of solvent in a specific region under influence of elasto-osmosis causes fluctuations in the local solute concentration, which may lead to *motility-induced osmo-elastic phase separation*.

We demonstrate the phenomenon in FIG 3, which presents 25 consecutive and temporally evenly spaced snapshots of a calculation considering an epithelium for which *χ*^(*α*)^ >> *X*^(*β*)^, i.e. *χ*^(*α*)^ = 10.5 and *X*^(*α*)^ = 2.5 (see Table S-I [26]), so of which the cells are fully permeable to the solvent but only marginally permeable to the solute. Cellular motion is predominantly deterministic on account of a low rotational diffusivity. Time progresses from the top left to the bottom right. The overall mean solute concentration is 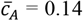, which due to partitioning results in a mean concentration per cell 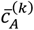 of roughly 0.15 to 0.17 at 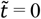, with 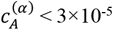. The individual panels show the solute volume fraction and the position of the membranes plotted on the same color.

**FIG 3.**
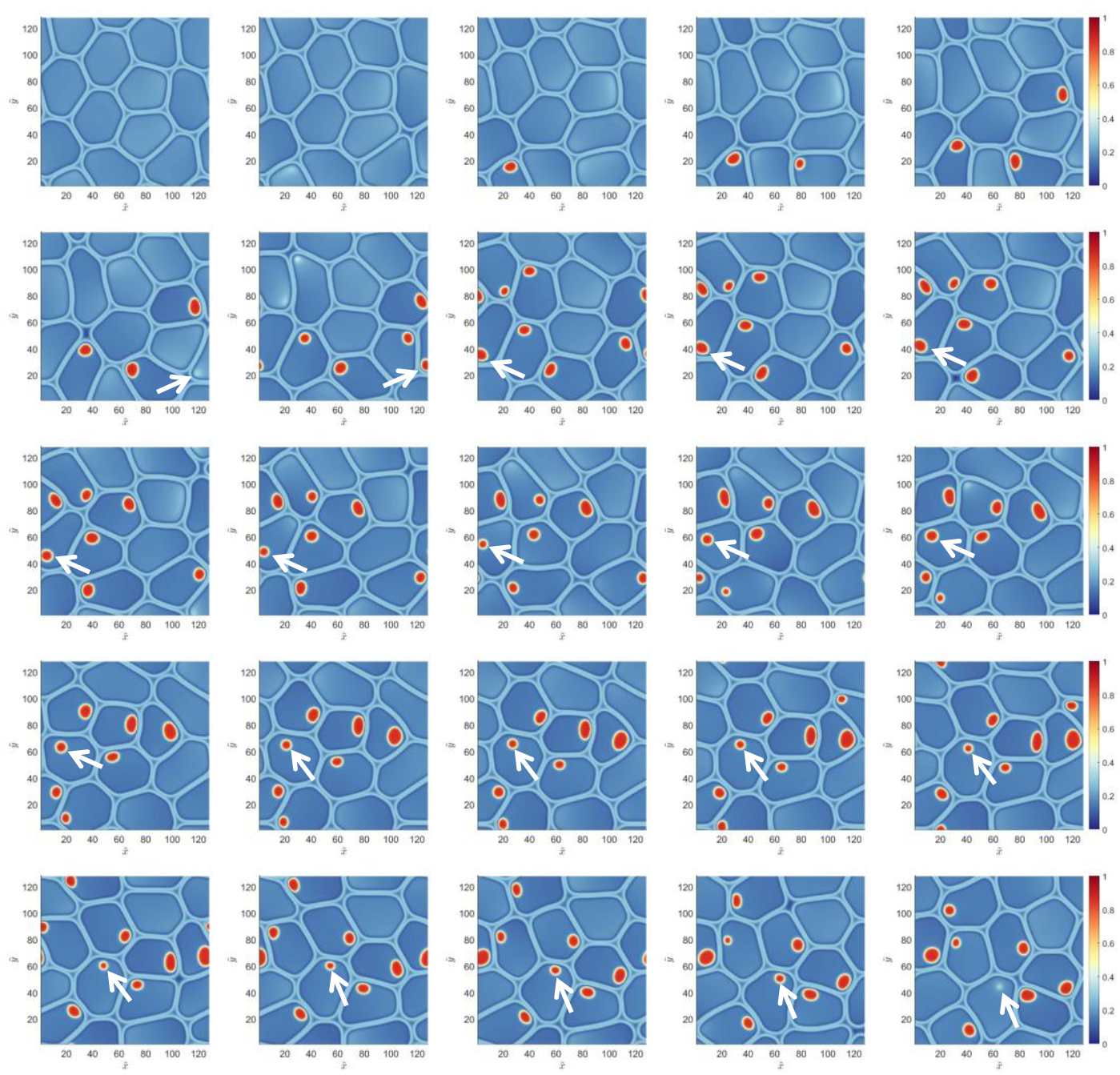
Dynamic numerical simulation of a confluent epithelium comprising twelve motile semipermeable cells (*n*= 12) containing a binary solution (*m*= 2), followed in time. All input parameters are given in Table S-I [26]. Time progresses in dimensionless units from the top left 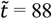 to 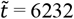 at the bottom right, out of a total integration time of 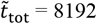. The color scale represents the local solute volume fraction, as well as the loci of the cell characterized by 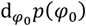 with 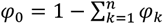 and scaled by a factor 1/6. Condensates show up in red. The white arrows track a condensate during its lifetime as its hosting cell migrates. This figure corresponds to *SupplementaryMovie1* [28].

As the cells migrate, fluctuations in concentration occur due to deformation, compression and extension. As these fluctuations locally cross the phase boundary, condensates form, change size and occasionally re-dissolve (follow the white arrows), depending on whether their hosting cell is deformed, compressed or expanded. In comparison to those encountered in real cells, the condensates in these calculations are unrealistically large. This is however deliberate. Exaggerating condensates size avoids numerical difficulties associated with the fact that the cell radius is two orders of much larger than that of a ‘typical’ membrane-less organelle, and show that motility and phase separation are *mutually* coupled: motility does not merely drive the formation and dissolution of condensates but, as we show below, the osmotic cell size modulation (discussed in FIG 2d) influences the fluidity of the epithelium. Although also likely present for smaller condensates, this effect is more pronounced for large condensates.

FIG 4 presents the analysis of the numerical simulation above, including the mean and maximum solute concentration, as well as the volume and center-of-mass (COM) displacement as a function of time for each cell individually. The black dash-dotted line in FIG 4a shows that the mean volume fraction remains at 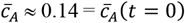, demonstrating that mass is near perfectly conserved. In contrast, the fluctuations in mean protein concentration *per cell* (colored lines) are non-negligible and seem to increase somewhat as time progresses. The fact that the fluctuations in cell volume are similar but inverted (FIG 4b) shows that variations in solute concentration are predominantly due to loss and gain of solvent upon compression and extension (thick lines, left axis. This ‘semi-permeable’ behavior is more clearly expressed by FIG 4c, where concentration (solid) and inverted volume (dashed) are presented in a scaled way. Clearly, the two properties generally coincide with some deviation where the solid actually to minor extent does traverse the cell membrane. FIG 4b shows that fluctuations in volume are still relatively small, remaining limited to below 10%.

**FIG 4.**
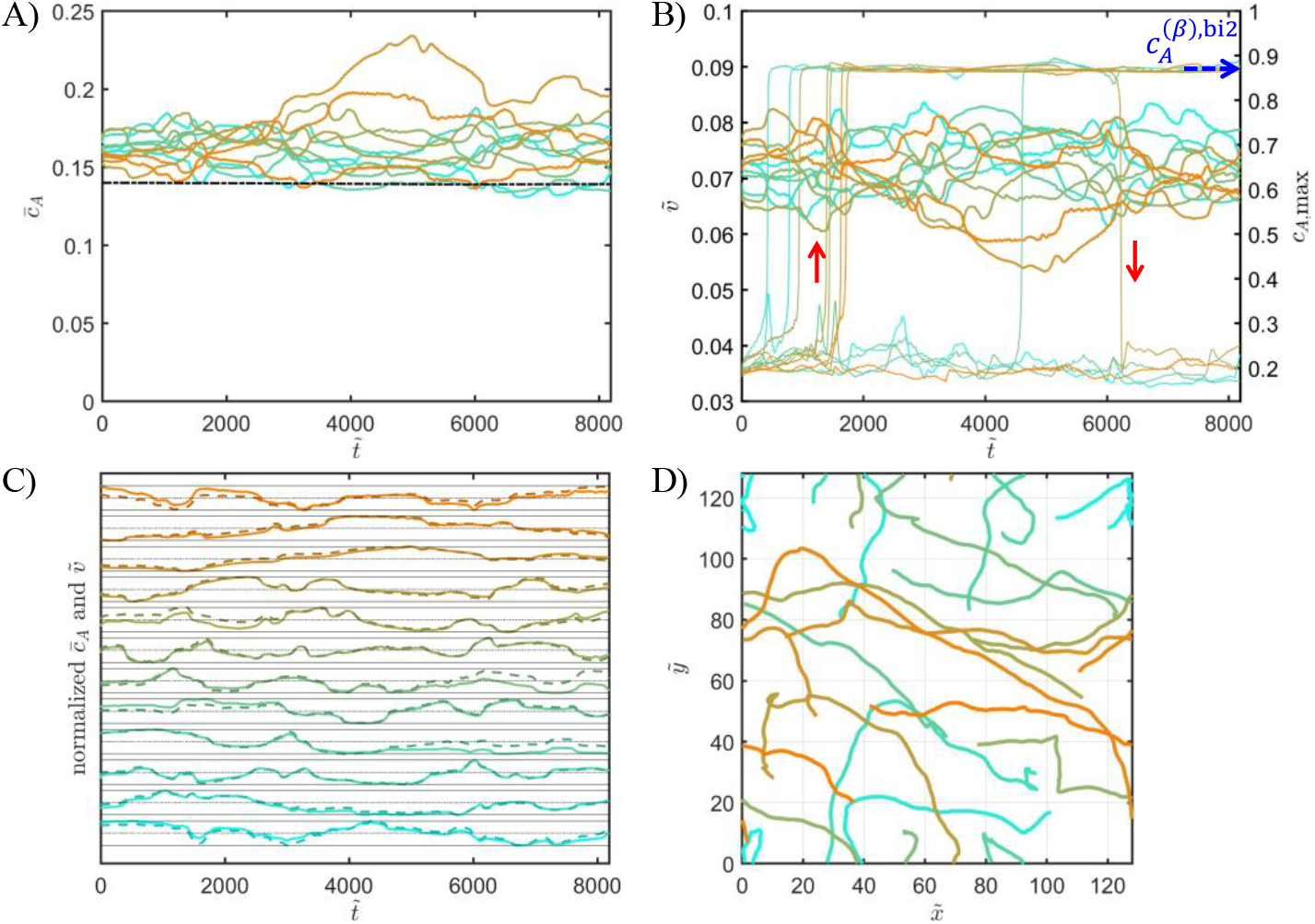
Analysis of the numerical simulation displayed in FIG 3. (a) overall mean solute volume fraction (dash-dotted black line) and mean volume fraction per cell (colored lines) plotted as a function of time. (b) Individual cell volumes (thick colored lines) and maximum solute volume fraction in each cell (thin colored lines), plotted as a function of time. (c) Mean solute concentration (solid) and the inverse of the cell volume (dashed), normalized by the respective maximum values and plotted as a function of time for each cell. (d) Centre-of-mass displacement of each cell. This figure corresponds to *SupplementaryMovie1* [28].

FIG 4b also reveals condensate formation and dissolution in individual cells as jumps in the maximum concentration (thin lines, right axis). For *χ*^(*β*)^ = 2.5, the condensate concentration is given by 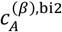 as indicated. The plot shows that a number of condensates form relatively early on, one of them redissolving around 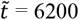 (follow red arrows on the brown curve) as the fluctuations in cell volume increase. This particular condensate corresponds to the one tracked by the white arrows in FIG 3. Finally, FIG 4d shows the COM-trajectories of the twelve cells during the numerical simulation. Clearly, for the present input, the epithelium has a high fluidity: cells migrate over long distances with mean square displacements reaching substantially beyond the typical cell radius (compare FIG 3). The fact that some of the trajectories run parallel expresses collective motion, typically observed both computationally [15] and experimentally [1].

To demonstrate the impact of the nature of cell motion on condensate formation, we next perform a set of numerical simulations at a stepwise increased rotational diffusivity. As mentioned, this parameter enters the stochastic motile component, which means that at elevated levels the cells and their contents become subject to fast, jerky motion, rather than smooth, long range spatiotemporally correlated migration as seen in the calculation described in FIG 3 and 4. The partitioning strength, *i.e*. set by *χ*^(*α*)^ and *χ*^(*β*)^, is left unchanged and so are all other input parameters (see Table S-I [26]). FIG 5 shows the analyses, together with the final snapshot of numerical simulations for *DR* = 1, 5 and 10. For *D*_*R*_ = 1, the epithelium is still fluid, evidenced by the fact that the COM-displacement is large. The hallmarks of a highly fluidity epithelium observed for *D*_*R*_ = 0.1, such as irregular cell shape, substantial fluctuations in solute concentration and volume, long range displacement and condensate formation, are retained for *D*_*R*_ = 1. Still, condensate formation becomes somewhat suppressed relative to *D*_*R*_ = 0.1 (compare FIG 5b with FIG 4b).

**FIG 5.**
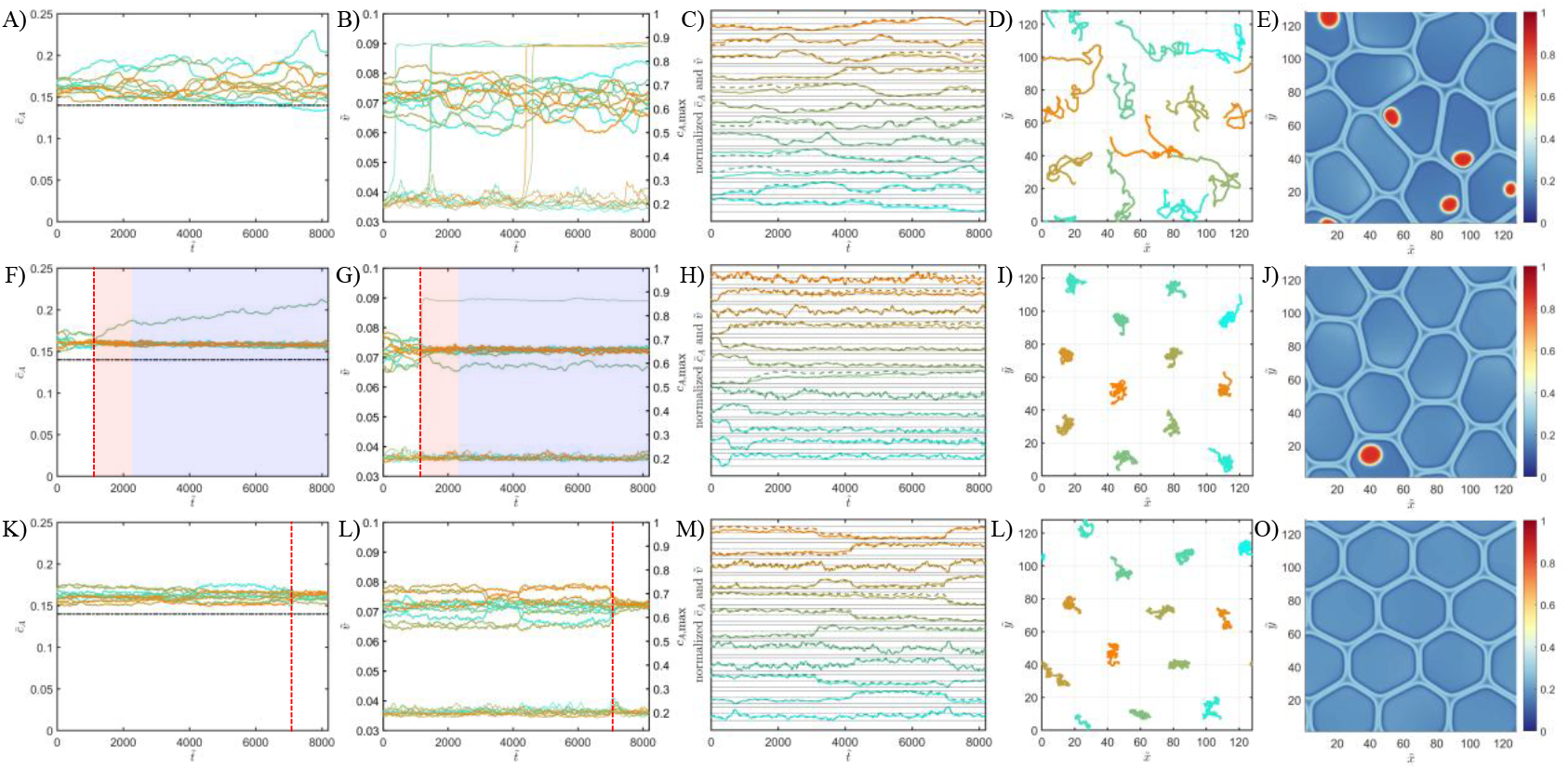
Numerical simulation of motile cells (*n*= 12) containing a binary solution (*m*= 2), as a function of cell rotational diffusivity. Left to right: *D*_*R*_ = 1, 5 and 10 in dimensionless units. All input parameters are given in Table S-I [26]. (a), (f), (k) mean solute concentration per cell (colored lines) and overall mean concentration (black dash-dotted lines), (b), (g), (l) fractional volume of each cell (thick lines, left axis) and maximum solute concentration in each cell (narrow lines, right axis), (c), (h), (m) mean solute concentration (solid) and the inverse of the cell volume (dashed), normalized by the respective maximum values, (d), (i), (n) center-of-mass displacement of each cell, (e), (j), (o) final snapshot of the numerical simulation; the color scale represents the local solute volume fraction, as well as the loci of the cell membranes as a function of the magnitude of the order parameter gradients. The columns in this figure correspond (left to right) to *SupplementaryMovie2, SupplementaryMovie3* and *SupplementaryMovie4* [28].

However, upon further enhancing stochastic motion, *i.e*. for *D*_*R*_ = 5 and *D*_*R*_ = 10, the behavior is markedly different. Displacement is short ranged (compare FIG 5d with 5i and 5n), indicating a transition to a low energy state in which the cells adopt a predominantly hexagonal arrangement ( I 5j and 5o, wherein they become ‘caged’ and wobble around their position in the lattice without exhibiting long range displacement. The transition to this energetically favorable ‘glassy’ or ‘crystalline’ state is relatively sharp and occurs at times 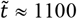 and 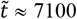 for *D*_*R*_ = 5 and *D*_*R*_ = 10, respectively (dashed red lines in FIG 5f, 5g, 5k and 5l). Interestingly, for *D*_*R*_ = 5 the transition occurs much sooner than for *D*_*R*_ = 10, likely because of a considerate deterministic contribution to the amplitude of spatial fluctuations in case of the former which now causes fast relaxation instead of maintaining the system far out of equilibrium.

FIG 5f, 5g, 5k and 5l show that following the transition to the hexagonal state, the mean solute concentrations and cell volumes more or less equalize, with fluctuations in both quantities becoming very small. As a result, for high values of *D*_*R*_ condensate formation is strongly suppressed. For *D*_*R*_ = 5 only one single condensate forms. Interestingly, its nucleation coincides with a fast relaxation of the epithelium to the hexagonal morphology. This, together with the initial decrease in the volume of this cell (FIG 5f and 5g, red shaded region), supports the possibility that the osmotic cell size depression accompanying the condensation temporarily increases the fluidity of the epithelium, thereby accelerating the morphological transition, which points to a feedback mechanism between motility and condensate formation. At later times, the mean solute concentration of the accommodating cell slowly increases with time, whereas its volume remains fairly constant (blue shaded regions). The accompanying *SupplementaryMovie3* [28] reveals that this concentration increase is due to slow growth of the condensate. In other words, owing to the fact that for the given *χ*^(*α*)^ and *χ*^(*β*)^ solute permeability is small but non-zero, the condensate collects material from neighboring cells to reduce free energy associated with its curvature.

In our final exploration, we demonstrate a second feedback mechanism that shows how the motility, or rather the fluidity, of the epithelium depends on the degree of ‘semipermeability’ set by the partitioning of the solute. In I 6, we compare motility in epithelia comprising ‘semipermeable’ (left column with ‘permeable’ cells (right column). The ‘semipermeable’/strong partitioning case is the same calculation as depicted in FIG 3 and 4 and for convenience repeated. For the ‘permeable’ case we impose a weak preference for the cytoplasm through *χ*^(*α*)^ = 4, rather than *χ*^(*α*)^ = 10, in fact as in FIG 2. All other input is left unchanged (see Table S1). In both cases, cell motion is dominantly deterministic, corresponding to a low value for the rotational diffusivity of *D*_*R*_ = 0.1.

Comparison between the left (‘semipermeable’ and right (‘permeable’ panels clearly shows that, despite equal kinetic parameters, whether or not the solute easily diffuses across cell boundaries drastically influences not only individual cell motility, but also the overall fluidity of the epithelium and, concomitantly, fluctuations in cell shape and volume. Reiterating our findings above: for strong partitioning and deterministic motion, anticorrelated fluctuation in mean concentration and cell volume lead to extensive condensation and large COM displacement, with the morphology staying out of equilibrium rather than adopting the energetically favored hexagonal arrangement (FIG 6a – 6e). In contrast, just by lowering *χ*^(*α*)^ to give weak partitioning, the cell membranes are effectively invisible to solute transport, which means that the volume and the mean concentration per cell are uncorrelated, as expressed by FIG 6f, 6g and 6h. As a result, the concentration barely fluctuates and despite the deterministic nature of the cell motion, now a transition to a stable hexagonal morphology takes place (at 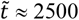, see FIG 6g and 6j) without any condensation taking place. Interestingly though, the ensemble is not jammed, as observed for strong partitioning in combination with highly stochastic motion (see FIG 5 middle and right column). Instead, as shown by FIG 6I and *SupplementaryMovie5* [28], cells keep migrating but in a strongly collective manner, which does not seem to break the hexagonal arrangement.

**FIG 6.**
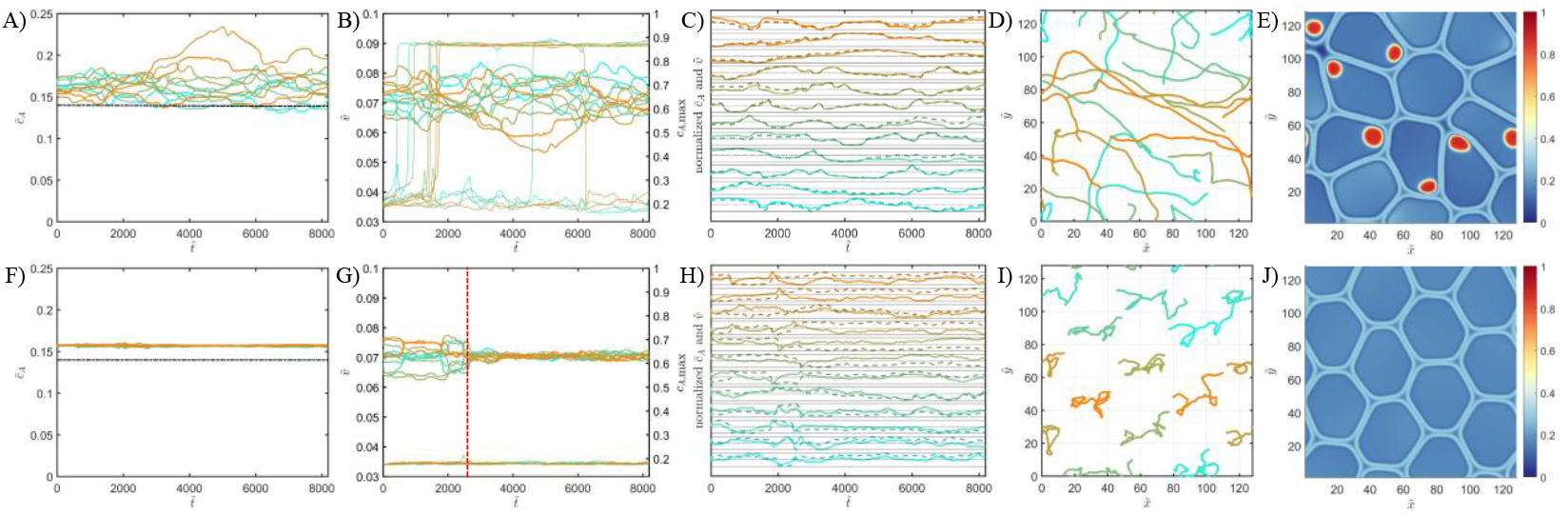
Numerical simulation of motile cells (*n*= 12) containing a binary solution (*m*= 2), as a function of the solute partitioning strength: *X*^(*α*)^ = 10.5 (left) and *X*^(*α*)^ = 4 (right), with *X*^(*β*)^ = 2.5 in both cases. All other input parameters are given in Table S-I [26]. (a) and (f) mean solute concentration per cell (colored lines) together with the overall mean concentration (black dash-dotted lines), (b) and (g) fractional volume of each cell (thick lines, left axis) together with the maximum solute concentration in each cell (narrow lines, right axis), (c) and (h) mean solute concentration (solid) and the inverse of the cell volume (dashed) normalized by the respective maximum values, (d) and (i) the center-of-mass displacement of each cell, (e) and (j) final snapshot of the numerical simulation; the color scale represents the local solute volume fraction, as well as the loci of the cell membranes as a function of the magnitude of the order parameter gradients. The vertical red dashed line in panel (g) indicates the point where the epithelium relaxes towards a predominantly hexagonal cell arrangement. The columns in this figure correspond (left to right) to *SupplementaryMovie1* and *SupplementaryMovie5* [28].

The numerical simulations in FIG 6 show that the partitioning strength directly influences the achievable range of the osmo-elastic potential Π_com_ during confluent cell migration for a given set of dynamic parameters, mean concentration and cell stiffness. We have shown above in FIG 2 that for the weak partitioning case of *χ*^(*α*)^ = 4 and *χ*^(*β*)^ = 2.5, for which we do *not* observe condensate formation, phase separation *is* principally possible owing to the double well nature of the free energy of the cytoplasm. The fact that it doesn’t occur in the numerical simulation represented by FIG 6f – 6j hence means that due to rather unrestricted solute transport across cell boundaries, Π_com_ remains too low to push the point of contact on the cytosol free energy of common tangent **4** near or beyond 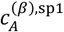, *i.e*. to sufficiently destabilize the solution (see FIG 2b). What this tells us is that condensate formation within a confluent epithelium, is not only determined by the solubility of a protein in the cytosol itself, but also by its compatibility with the extracellular environment.

## IV. DISCUSSION

This work computationally explores how changes in volume and shape of motile cells in a confluent epithelial tissue give relate to the formation and dissolution of cytoplasmatic biocondensates. Our calculations reveal an intricate, so far unexplored interplay between osmosis, cell bulk elasticity, individual cell motility and global epithelial fluidity. In the first part of the work, we study the phase behavior of a solute partitioning across membrane of a single cell, as a function of the pressure experienced by the cell, either through external force (out of equilibrium) or through a balancing of the osmotic pressure by the cell’s own bulk stiffness (equilibrium. The latter yields an expression for the ‘osmo-elastic’ potential, linking the cell’s stiffness and size to the miscibility of the solute. Our calculations reveal stable, metastable and unstable regimes in a phase diagram mapped as a function of solute concentration and the (applied) potential and predict a change in the cell size when nucleating a condensate in a metastable cytoplasm.

In the second part, we explore out-of-equilibrium motility in the epithelial tissue by numerically integrating the coupled evolution equations for the field parameters that describe the cells and the solute concentration. Strong partitioning in favor of the cytoplasm results in an effective ‘semipermeability’, which prevents transport of solute across cell membranes. Consequently, motility-induced fluctuations in cell shape and size give rise to local variations in concentration, as well as condensate formation and dissolution, in particular if cell motion is deterministic. Such a ‘semipermeable’ epithelium has a high fluidity with cells exhibiting large center-of-mass displacement. If cell motion becomes predominantly stochastic, the tissue relaxes to an energetically favored ‘crystalline’ state in which the cells adopt a hexagonal arrangement. In this case, cell volumes equalize and condensate formation is suppressed.

Condensate formation is also suppressed if the partitioning strength is low. Now cells become ‘permeable’ and ‘invisible’ to solute transport, which is no longer subject to osmo-elastic fluctuations. Interestingly, in a feedback mechanism, even if cell motion is predominantly deterministic, the absence of condensation strongly reduces individual displacement and encourages relaxation to a hexagonal arrangement which remains in-tact despite extensive collective motion. In other words, our calculations show that, motility in a tissue does not merely induce condensation and dissolution, but the latter in turn affect not only individual cell motility, but also the fluidity of the epithelium as a whole.

Coupling phase behavior to cell motility produces a very rich behavior, of which we feel we have so far only scratched the surface. In its current state, we expect our model to reveal further interesting phenomena, already through straightforward exploration of the parameters space, *e.g*. by varying the number of components, the cell stiffness and interaction. Nevertheless, we acknowledge that the present model may perhaps be best considered ‘bio-inspired’, rather than ‘biological’ and would certainly need experimental validation. Integrating features essential to motility in biological tissue, such as actin and myosin action, cell-cell and cell-substrate adhesion, as well as non-equilibrium processes will be focus of future work. Although we did not set out to solidly prove that motility-induced condensate formation occurs in life cells, our observations certainly fit very well with the notion that i) the dynamic and mechanical behavior of a tissue results from a superposition of long-range phenomena that emerge from collective local (sub)cellular processes and ii) that biological function and self-regulation critically relies on feedback mechanisms through mutually dependent processes.

To end on an interesting fundamental note, if condensation-induced cell size modulation indeed affects the fluidity of the epithelium upon breaking the hexagonal symmetry, the jammed state may hence ‘melt’ on account on solute concentration or solvent interaction. Interestingly, if the solute-solvent phase diagram has an UCST, melting would occur by *lowering* the temperature.

## V. CONCLUSIONS

Using biologically numerical simulations, we show that cell motility and cytoplasmatic phase behavior in confluent epithelial tissues may well be coupled phenomena. For this, we have developed a multiphase-multicomponent model that integrates a typical order-parameter based free energy for a confluent 2D tissue with a Flory-Huggins based mixing contribution that takes any number of solutes. Interaction parameters interpolate smoothly between extremes defined for the cytoplasm and the extracellular environment, either globally or for each cell individually. Elastic pressure modulations in cell size and shape induced by migration trigger phase separation of a solute (protein) if its concentration is near its saturation point. Condensate formation depends on the partitioning strength of a solute between the extracellular environment and the cytoplasm, as well as on whether cell motion is predominantly deterministic or stochastic. Interestingly, osmotic changes in cell size associated with phase separation seem, in turn, to affect cell motility and even the fluidity of the full epithelium.

## References

[1] B. R. Brückner and A. Janshof, Importance of integrity of cell cell junctions for the mechanics of confluent MDCK II cells, Sci. Rep. 8, 141117 (2018).

[2] D. M. Hatters, Grand challenges in biomolecular condensates: structure, function, and formation, Front. Biophys. 1, 1208763 (2023).

[3] Y. Li, Y. Liu, X. Y. Yu, Y. Xu, X. Pan, Y. Sun, Y. Wang, Y. H. Song and Z. Shen, Membraneless organelles in health and disease: exploring the molecular basis, physiological roles and pathological implications, STTT 9, 305 (2024).

[4] K. Graham, A. Chandrasekaran, L. Wang, A. Ladak, E. M. Lafer, P. Rangamani, Liquid-like VASP condensates drive actin polymerization and dynamic bundling, Nat. Phys. 19, 574 (2023).

[5] T. Wiegand and A. A. Hyman, Drops and fibers — how biomolecular condensates and cytoskeletal filaments influence each other, Emerg. Top. Life Sci. 4, 247 (2020).

[6] D. Sun, I. LuValle-Burke, K. Pombo-García and A. Honigmann, Biomolecular condensates in epithelial junctions, Curr. Opinion in Cell Biol. 77, 102089 (2022).

[7] D. Sun, X. Zhao, T. Wiegand, C. Martin-Lemaitre, T. Borianne, L. Kleinschmidt, S. W. Grill, A. A. Hyman, C. Weber and A. Honigmann, Assembly of tight junction belts by ZO1 surface condensation and local actin polymerization, Dev. Cell 60, 1234 (2025).

[8] Y. Wang, C. Zhang, W. Yang, S. Shao, X. Xu, Y. Sun, P. Li, L. Liang and C. Wu, LIMD1 phase separation contributes to cellular mechanics and durotaxis by regulating focal adhesion dynamics in response to force, Dev. Cell. 56, 1313 (2021).

[9] S. Banjade and M. K. Rosen, Phase transitions of multivalent proteins can promote clustering of membrane receptors, eLife 3, e04123 (2014).

[10] S. Jeon, Y. Jeon, J. Y. Lim, Y. Kim, B. Cha and W. Kim, Emerging regulatory mechanisms and functions of biomolecular condensates: implications for therapeutic targets, STTT 10, 4 (2025).

[11] D. Ngaia, A. L. Mohabeer, A. Mao, M. Lino and M. P. Bendeck, Stiffness-responsive feedback autoregulation of DDR1 expression is mediated by a DDR1-YAP/TAZ axis, Matrix Biol. 110, 129 (2022).

[12] S. Torrino, W. M. Oldham, A. R. Tejedor, I. S. Burgos, L. Nasr, N. Rachedi, K. Fraissard, C. Chauvet, C. Sbai, B. P. O’Hara, Mechano-dependent sorbitol accumulation supports biomolecular condensate, Cell 188, 447 (2025).

[13] N. Sanfeliu-Cerdan and M. Krieg, The mechanobiology of biomolecular condensates, Biophys. Rev. 6, 011310 (2025).

[14] L. M. Faure, V. Venturini and P. Roca-Cusachs, Cell compression – relevance, mechanotransduction mechanisms and tools, J. Cell Sci. 138, jcs263704 (2025).

[15] B. A. Camley and W.-J. Rappel, Physical models of collective cell motility: from cell to tissue, J. Phys. D Appl. Phys. 50, 113002 (2017).

[16] D. Wenzel and A. Voigt, Multiphase field models for collective cell migration, Phys. Rev. E 104, 054410 (2021).

[17] S. Monfared, A. Ardaševa1 and A. Doostmohammadi, Multiphase-Field Models of Tissues, Annu. Rev. Condens. Matter Phys. 17, 91 (2026).

[18] S. Seirin Lee, S. Tashiro, A. Awazu and R. Kobayashi, A new application of the phase-field method for understanding the mechanisms of nuclear architecture reorganization, J. Math. Biol. 74, 333 (2017).

[19] D. M. Saylor, C.-S. Kim, D. V. Patwardhan and J. A. Warren, Diffuse-interface theory for structure formation and release behavior in controlled drug release systems, Acta Biomater. 3, 851 (2007).

[20] J. J. Michels, K. Zhang, P. Wucher, P. M. Beaujuge, W. Pisula1,3 and T. Marszalek, Predictive modelling of structure formation in semiconductor films produced by meniscus-guided coating, Nat. Mater. 20, 68 (2021).

[21] G. Zhang, R. Mueller, A. Doostmohammadi and J. M. Yeomans, Active inter-cellular forces in collective cell motility, J. R. Soc. Interface 17, 20200312 (2020).

[22] P. C. Hohenberg and B. I. Halperin, Theory of dynamic critical phenomena, Rev. Mod. Phys. 49, 435 (1977).

[23] M. Chiang, A. Hopkins, B. Loewe, D. Marenduzzo, and M. C. Marchetti, Multiphase field model of cells on a substrate: From three dimensional to two dimensional, Phys. Rev. E 110, 044403 (2024).

[24] S. Henkes, K. Kostanjevec, J. M. Collinson, R. Sknepnek, E. Bertin, Dense active matter model of motion patterns in confluent cell monolayers, Nat. Commun. 16, 1405 (2020).

[25] C. Schaefer, J. J. Michels, P. van der Schoot, Structuring of thin-film polymer mixtures upon solvent evaporation, Macromolecules 49, 6858 (2016).

[26] See Supplemental Material [url] for details on calculations and input parameters.

[27] J. Fritzen, A. Samanta, N. S. Kuhr, A. Preuß, E. L. Sternburg, L. S. Stelzl, J. J. Michels, D. Dormann and A. Walther, biorxiv, 10.64898/2026.04.07.716875.

[28] See Supplemental Material [url] containing dynamic animations of the numerical simulations of confluent cell motility.

