## Supplementary Information for "Elasto-Osmotic Phase Separation in Confluent Cellular Tissues"

*Jasper J. Michels**

Max Planck Institute for Polymer Research, Ackermannweg 10, 55128 Mainz, Germany

SUPPLEMENTARY INFORMATION

### Section S-I. Calculation and model input

All dynamic calculations are performed on an $N$-sized square grid with periodic boundary conditions, using a straightforward forward-time-central-space finite difference algorithm. In Table S-I we list all input parameters for the calculations in this work, while referring to the main text figures where appropriate. If no reference is made, the value is left unchanged in all calculations. For convenience, all quantities are assumed dimensionless. We nevertheless provide the original dimension for clarity.

**Table S-I.** Model input parameters used in the calculations described in the main document

| parameter | Description | original units | non-dimensional value |
|---|---|---|---|
| $a$ | fundamental length scale | m | 0.01 |
| $A$ | Landau prefactor | J $m^{-3}$ | 1.0 |
| $B$ | cell 'stiffness' | J | 100 |
| $\lambda^2$ | order parameter gradient energy coefficient | J $m^{-1}$ | 1.5 |
| $\kappa_{ii}$ | concentration gradient energy coefficient | J $m^{-1}$ | 0.15 |
| $D$ | self-diffusivity | $m^2$ $s^{-1}$ | 1.0 |
| $D_R$ | rotational diffusivity | $rad^2$ $s^{-1}$ | 0.1[a]; 1.0[b]; 5.0[c]; 10[d] |
| $\omega$ | cell-cell overlap penalty | J $m^{-3}$ | 20 |
| $\Gamma$ | conversion rate | $s^{-1}$ | 1.0 |
| $\alpha$ | active force magnitude | N | 0.005 |
| $\xi$ | friction coefficient | kg $s^{-1}$ | 0.125 |
| $\bar{c}_A$ | mean solute volume fraction | - | 0.14 |
| $\chi^{(\alpha)}$ | solute-solvent interaction parameter in the ECE | - | 3[e]; 4[f]; 10.5[g] |
| $\chi^{(\beta)}$ | solute-solvent interaction parameter in the ECE | - | 1.5[h]; 2.5[i] |
| $N$ | numerical grid size | - | 128 |

| | | | |
|---|---|---|---|
| $\Delta\tilde{t}$ | dimensionless time step | - | 0.0025 |
| $\tilde{t}_{\mathrm{tot}}$ | total integration time | - | 8192 |

[a]Figure 3, 4, 6 (left column), S-I; [b]Figure 5 (left column); [c]Figure 5 (middle column); [d]Figure 5 (right column); [e]Figure 1; [f]Figure 2, 6 (right column); [g]Figure 3, 4, 5, 6 (left column), S-I; [h]Figure 1; [i]Figure 2, 3, 4, 5, 6, S-I.

### Section S-II. Mass conservation reference calculation

As explained in the main text, we apply a Helmholtz decomposition to the divergent total velocity field resulting from cell migration. Prior to decomposition we smoothen the velocity field using an 11x11 grid cell block smoothing procedure. Main text Figures 4 – 6 (in particular the dash-dotted block lines in the topmost panels) show that coupling of the solute concentration to the solenoidal part of the smoothed total cellular velocity preserves mass very effectively. As a reference, Figure S-I presents the same calculation as presented in main text Figures 3 and 4, without applying the Helmholtz decomposition. In particular Figure S-Ia and S-Ib clearly show that now mass is not conserved as the overall and individual mean concentrations, as well as the maximum concentrations per cell, decrease over time. As a result, the solution in the cytoplasm stabilizes and no phase separation occurs (see Figure S-If).

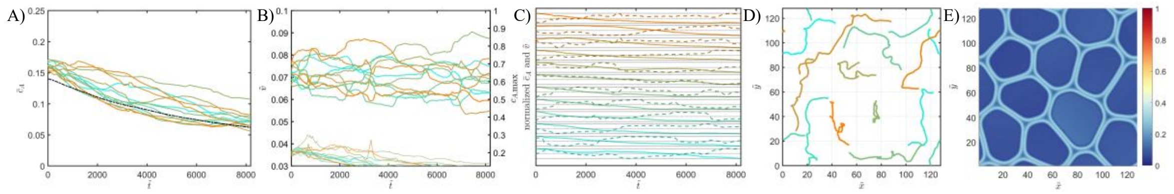


**Figure S-I.** *Numerical simulation of semipermeable motile cells, without performing a Helmholtz decomposition of the cell-associated velocity field. Input parameters are given in Table S-I. Panels (a) – (e) respectively show the mean solute concentration per cell (colored lines), as well as the overall mean concentration (black dash-dotted lines), the fractional volume of each cell (thick lines, left axis) and the maximum solute concentration in each cell*

*(narrow lines, right axis), the mean solute concentration (solid) and the inverse of the cell volume (dashed), normalized by the respective maximum values, the center-of-mass displacement of each cell and the final snapshot of the numerical simulation; the color scale represents the local solute volume fraction, as well as the loci of the cell membranes as a function of the magnitude of the order parameter gradients. The figure corresponds to SupplementaryMovie6.*
